# Clonality-Related Traits Add Independent Specialization Axes to Herbs’ Trait Strategies

**DOI:** 10.1101/2023.03.15.532195

**Authors:** Stefano Chelli, Jitka Klimešová, James Lee Tsakalos, Giacomo Puglielli

**Affiliations:** School of Biosciences and Veterinary Medicine, Plant Diversity and Ecosystems Management Unit, University of Camerino, Camerino, Italy; Department of Experimental and Functional Morphology, Institute of Botany of the Czech Academy of Sciences, Třeboň, Czech Republic; Department of Botany, Faculty of Science, Charles University, Praha, Czech Republic; Harry Butler Institute, Murdoch University, Murdoch, Perth, WA, Australia; Departamento de Biología Vegetal y Ecología, Facultad de Biología, Universidad de Sevilla, Apartado 1095, 41080– Sevilla, Spain

## Abstract

The functional diversity of vascular plants is remarkable. Yet, previous studies showed that trait trade-offs constrain aboveground or fine-root trait variation. How do neglected functions such as resprouting and clonal growth, key for fitness maintenance in some plant groups, integrate in these trait frameworks? By using an extensive dataset (> 2000 species) spanning aboveground, fine-root and clonality-related traits of herbs, we asked whether clonal traits relate to species positioning in the aboveground or fine-root trait spaces. Clonal and non-clonal herbs were undistinguishable in the aboveground or fine-root trait spaces. Clonality-related traits were also weakly coordinated with the other trait dimensions. Altogether, these results suggest that clonality-related traits add independent functional specialization axes to plants’ trait strategies. We identified two potential specialization axes in clonal traits. The first axis summarizes the positive scaling between bud bank size and persistence of clonal connections, reflecting species specialization for on-spot persistence and tolerance to disturbance (*persistence* axis). The second axis, summarizes the positive scaling between multiplication rate and lateral spread, reflecting specialization for clonal multiplication and acquiring new space in horizontal dimension (*clonal multiplication* axis). We call for integrating these axes in existing strategy schemes to fully elucidate the multidimensional trait strategies of plants.

## Introduction

Plants continuously acquire, use and store above- (i.e., Carbon) and belowground (i.e., water and nutrients) resources to serve multiple functions related to growth, survival, and reproduction (Weiher et al., 1999). Different functional traits (*sensu* Violle et al. 2007) have been successfully used as proxies of these functions. Thus, trait combinations are now used to describe the life history strategies employed by plants to maximize fitness across a range of environmental conditions, and across the tree of life (e.g., Westoby et al. 2002; Reich et al. 1997). This trait-based approach (*sensu* Westoby & Wright 2006) continues to identify leading and often independent trait dimensions (e.g., leaf economics and size-related traits; Diaz et al. 2016), each summarizing major trait trade-offs (e.g., the leaf economics spectrum; Wright et al. 2004), and these dimensions reflect evolutionary and energetic constraints over functional differentiation in all major groups of vascular plants. However, how many independent dimensions of ecological specialization are needed to describe plant strategies is yet to be determined.

At the global scale, functional differentiation among vascular plants is summarized by two independent trait dimensions defining the so-called Global Spectrum of Plant Form and Function (Díaz et al. 2016; referred to as aboveground trait space). This trait space is defined by six traits - plant height, specific stem density, seed mass, individual leaf size, specific leaf area, and leaf nitrogen content on a mass basis - that are known to reflect key mechanical and energetic trade-offs underlying the aboveground vascular plant trait strategies (**Table S1**; Díaz et al. 2016; but see also Westoby et al. 2002). The aboveground trait space summarizes how the independent plant economics and organ size-related trait dimensions define plant resource acquisition and use strategies that are related to species’ growth, survival, and reproduction. Since its formulation, the aboveground trait space represents a leading trait-framework for describing and comparing aboveground vascular plant functional strategies across all levels of biological organization and spatial scales (Bruelheide et al., 2018; Thomas et al., 2020; Joswig et al., 2022).

The description of the aboveground trait space triggered interest about how belowground acquisitive organs can be integrated in this pattern. Recently, Carmona et al. (2021) combined global data for belowground fine-root economics traits (**Table S1**: specific root length, root tissue density, root nitrogen content and root diameter) to define the fine-root economics space. Finally, they integrated the root economics space in the aboveground trait space. Unlike previous interpretations (see, e.g., Reich 2014; Bergmann et al. 2021), which suggested root traits to reflect the same key trait trade-offs of their aboveground counterparts, Carmona et al. (2021) demonstrated that the aboveground and fine-root trait dimensions are orthogonal and, therefore, independent. Indeed, multiple studies (e.g., Weigelt et al. 2021; E-Vojtkó et al. 2022) have suggested that the possible number of trait dimensions summarizing the large variability of plants’ life-history strategies might be larger than expected on the basis of a single whole-plant economics spectrum (*sensu* Reich 2014) governing both aboveground and fine-root trait combinations.

Although the above-mentioned trait spaces covered many functional dimensions of vascular plants, there are still overlooked key functions that could be integrated in such frameworks – e.g., on spot persistence, space occupancy, resource sharing and storage, and clonal multiplication (Ottaviani et al. 2017; Klimešová et al. 2018). These functions are exerted by organs mostly located belowground - e.g., rhizomes, stolons, tubers, bulbs, etc. (Klimešová et al. 2019). For instance, among perennial herbs, such organs can be found in approximately 60% of European species (see Van Drunen & Husband 2019). Many perennial herbs can in fact rely on these organs to reproduce clonally as an alternative means to sexual reproduction (Aarssen 2008). In addition, clonal reproduction for these plant groups is essential for their spatial organization within communities, as well as for their competitive ability in horizontal dimension and resilience after biomass removal (e.g., after grazing, mowing, and fire; Pausas et al. 2018). All these functions are mostly modulated by clonal and bud-bank traits (Klimešová et al. 2017). Hence, in perennial clonal herbs, such traits are key to guarantee fitness maintenance and ultimately long-term population persistence under given environmental conditions (e.g., Mogie & Hutchings 1990). For this reason, clonal and clonality-related traits such as bud-bank traits, are considered one of the most prominent axes of variation in ecological strategies of many perennial herbs (Aarssen 2008; Klimešová et al. 2017).

However, it remains unclear how clonality integrates with other major axes of ecological differentiation in perennial herbs. Studies relating clonal and bud bank traits with the traits defining the aboveground trait space are sparse and mainly limited to specific leaf area, plant height, and seed mass (i.e., the traits defining the LHS scheme; Westoby 1998). Overall, these studies found that LHS traits were largely independent of clonal and bud bank traits in herbs (Herben et al. 2012; Herben et al. 2016; Klimešová et al. 2016; Herben et al. 2018). Only in some cases (e.g., see Klimešová et al. 2016; Lubbe et al. 2021), or when focusing on reduced species sets within a given habitat type (e.g., temperate grasslands; Ladouceur et al. 2019), some patterns emerge. Although organs other than roots can add complexity in terms of functions to the belowground trait space (Lachaise et al. 2022), the relationship between clonality-related traits with traits belonging to root economics is largely unknown (but see Klimešová & Herben 2023). Integrating clonal and bud bank traits of clonal perennial herbs within the independent aboveground and fine-root trait spaces is therefore a priority to potentially uncover overlooked aspects of the ecological strategies of this plant group.

Here, we explore how clonal and bud bank traits integrate within the aboveground and fine-root trait spaces of perennial clonal herbs. To do that, we combined the most extensive and curated dataset of plant clonal and bud-bank traits to date, the CloPla database (Klimešová et al. 2017), with the aboveground and fine-root trait frameworks provided by the aboveground (Díaz et al. 2016) and by the fine-root (Carmona et al. 2021) trait space. As a first step, we analyzed the extent by which perennial clonal herbs differ in terms aboveground and fine-root trait combinations from other non-clonal growth forms in our dataset (i.e., non-clonal woody, perennial non-clonal herbs and annual herbs). Similar trait combinations with other growth forms (most likely with other herbs) would suggest that being clonal does not necessarily correspond to peculiar trait syndromes, hinting that clonality might add a function that is not captured by the aboveground and fine-root trait combinations alone. Secondly, we explore the multivariate relationships between clonal and bud bank traits with the aboveground and fine-root trait combinations. In particular, we analyzed how single clonal and bud bank traits, as well as their combinations in a multivariate space, relate to aboveground and fine-root trait combinations for perennial clonal herbs. Altogether, our results show that clonality adds an independent function to the ones currently summarized by the aboveground and fine-root trait space frameworks.

## Methods Dataset

### Clonal and bud-bank traits

Clonal and bud bank traits were obtained from the CloPla database (Klimešová et al. 2017). CloPla contains data for 2909 clonal and non-clonal vascular plant species that are common across Central Europe. The dataset spans woody/non-woody, annual/non-annual plants, and clonality-related trait information. For our analysis, we selected the following traits (**Table S1**): (i) The *total bud bank size*, expressed as the total number of buds including stem buds at the soil level and root buds at different depths. (ii) The *multiplication rate*, calculated as the number of offspring shoots/parent shoot. (iii) The *lateral spread*, defined as the lateral spreading distance of clonal growth organs. (iv) The *persistence of clonal connection*, expressed as the lifespan of connection between offspring rooting units and parental rooting unit. We focused our analysis on these clonal traits because: (a) they reflect meaningful aspects of clonal plants ecological strategies along environmental gradients (e.g., Herben et al. 2018; Chelli et al. 2019; **Table S1**); (b) they have been recently included in a robust conceptual framework with the availability of standard protocols for field measurements (Klimešová et al. 2019); (c) they synthetize the main functions provided by the wide array of clonal and bud bank traits (see Introduction and Ottaviani et al. 2017).

### Aboveground and fine-root functional trait data

CloPla was combined with the aboveground and the fine-root economics functional traits information obtained from Carmona et al. (2021) (**Table S1**). Before merging the datasets, taxonomic information was homogenized following The Plant List v.1.1 using the R package ‘Taxonstand’ (Cayuela et al. 2012). Aboveground traits included: plant height (ph), specific stem density (ssd), seed mass (sm), individual leaf size (la), specific leaf area (sla), and leaf nitrogen content on a mass basis (ln). Trait information was available for 2674 species also included in CloPla (completeness: ph = 97%, ssd = 66%, sm = 78%, la = 72%, sla = 70%, ln = 43%). Fine-root economics traits included: specific root length (SRL), root tissue density (RTD), root nitrogen content (N) and root diameter (D). This trait information was available only for 474 species also included in CloPla (completeness: SRL = 82%, RTD = 68%, N = 50%, D = 83%). Due to the dramatic difference in the number of species between the aboveground and fine-root datasets, they were analyzed separately.

## Data analysis

### Trait space occupation by growth form

We first defined the aboveground and fine-root trait spaces using Principal Component Analysis (PCA) on imputed trait data by using the funspace function in the ‘funspace’ R package (Carmona et al. 2023) (see **Appendix S1; Fig. S1-S4; Table S2** for details on PCA and imputation procedure testing). To test how clonal plants differ in terms of aboveground and fine-root trait combinations compared to other non-clonal growth forms, we classified the species in our dataset as perennial clonal herbs (n = 1323 and 195, in the above- and belowground dataset), annuals (577 and 90), perennial non-clonal herbs (603 and 79), and non-clonal woody plants (152 and 66) using all the metadata available in CloPla (**Fig. S5-S6**). We then analyzed the extent of the shared portion of the trait space among growth forms using overlap-based dissimilarity (Carmona et al. 2016) quantified using kernel-density-based functions included in the TPD R package (Carmona et al. 2019). However, considering the different sample size among growth forms both in the aboveground and fine-root trait datasets, we assessed the overlap-based dissimilarity by using an equally sized random sample of species in each group (n = 50 for each growth form) across 500 iterations. PERMANOVA was also used to quantify how much of the trait variation along PC1 and PC2 of the aboveground or of the fine-root trait space was explained by the growth form.

### Linking clonal and bud bank traits to aboveground and fine-root trait combinations

To analyze how the aboveground and fine-root trait combinations are linked to clonality-related traits, we mapped the clonal and bud-bank related traits within the trait spaces defined by the aboveground and fine-root traits. For this analysis, since clonality-related traits were mostly available only for clonal plants, we redefined the aboveground and fine-root trait spaces using clonal plants only (i.e., perennial clonal herbs) (**Fig. S7-S8**). Our final datasets for this analysis included information for 1257 and 195 species for the aboveground and the fine-root dataset, respectively. Generalized Additive Models (GAMs) with a bivariate smoother were used to draw patterns of variation of clonal traits within the aboveground and fine-root traits space of clonal plants. GAMs were run using the funspaceGAM function included in the ‘funspace’ R package (Carmona et al. 2023) by setting a clonal trait as the response variable, and the trait space axes defining either the aboveground or the fine-root trait space of clonal plants as the bivariate explanatory variable. Finally, to test the consistency of the target relationships within functionally and ecologically distinct groups, clonal species were classified as ferns, forbs, and graminoids. However, ferns (n = 68) were only represented in the aboveground dataset and fine-root traits were missing for them. To seek comparability between aboveground and fine-root analyses, we restricted our analysis to forbs and graminoids (**Appendix S2; Fig. S9-S11**). The aboveground dataset included 919 forbs and 338 graminoids. The fine-root dataset included 110 forbs and 85 graminoids. For this test, we run GAMs and mapped their predictions only within the specific regions of the trait space occupied by each functional group (**Fig. S12-S19**).

To further test the relationship between clonal traits and aboveground and fine-root trait combinations, we used the following approach: (i) we characterized the clonal trait space using the species with complete trait information for total bud-bank size, multiplication rate, lateral spread, and persistence of clonal connections (n = 1181 species) via PCA. (ii) We tested for the correlation between the clonal trait space and its aboveground and fine-root counterparts using Procrustes analysis run via ‘protest’ function included in the vegan R package (Dray & Dufour 2007). This analysis rotates one configuration (the observed clonal trait space) to maximum similarity with a target one (either aboveground or fine-root trait space) and tests the non-randomness of the similarity via permutation (999 randomizations) based on Monte-Carlo simulations. All analyses were performed in R 4.2.0 (R Core Team, 2022).

## Results

### Trait space occupation by growth form

In terms of aboveground trait combinations, as expected, woody species were clearly different from the other growth forms (not-shared portion of the trait space = 0.48 ± 0.08), while perennial clonal and non-clonal herbs were very similar (overlap = 0.89 ± 0.07; **Table 1; Fig. S5**). In the fine-root trait space, differences between growth forms were less evident. However, perennial clonal and non-clonal herbs had the lowest dissimilarity (**Table 1; Fig. S6**). Overall, in the aboveground trait space, the main differentiation between growth forms was along PC2 (i.e., the plant and organ size axis) (**Fig. S5**), reflecting the difference between herbaceous and woody plants (**Fig. S5**). Accordingly, PERMANOVA analysis showed that growth form explained 6% and 31% of traits variation along PC1 and PC2 of the aboveground trait space. Concerning the fine-root trait space, the main differentiation among growth forms was observed along the specific root length – root diameter (SRL-D) axis (PC1; **Fig. S6**), but growth form explained only 7.5% and 3.2% of traits variation along PC1 and PC2 of the fine-root trait space, respectively.

**Table 1.** Overlap-based dissimilarity and its components (proportion of overlapping and non-overlapping portions of the trait space) among growth forms (A = annuals; PC = perennial clonal herbs; PNC = perennial non-clonal herbs; W = non-clonal woody plants) in the aboveground and fine-root trait spaces. Overlap-based dissimilarity was always evaluated at common sample size (n = 50 per each growth form) and the reported values are means and standard deviations (in parenthesis) calculated across 500 repetitions.

| **Aboveground space** |  |  |  |  |  |  |  |  |  |
| --- | --- | --- | --- | --- | --- | --- | --- | --- | --- |
|  | **Dissimilarity** |  |  | **Overlap** |  |  | **Not shared** |  |  |
|  | **PC** | **PNC** | **W** | **PC** | **PNC** | **W** | **PC** | **PNC** | **W** |
| **A** | 0.33 (0.06) | 0.31 (0.05) | 0.84 (0.04) | 0.86 (0.08) | 0.88 (0.07) | 0.40 (0.10) | 0.14 (0.08) | 0.12 (0.07) | 0.60 (0.10) |
| **PC** |  | 0.26 (0.05) | 0.75 (0.05) |  | 0.89 (0.07) | 0.56 (0.09) |  | 0.11 (0.07) | 0.44 (0.09) |
| **PNC** |  |  | 0.80 (0.04) |  |  | 0.55 (0.11) |  |  | 0.45 (0.11) |

| **Fine-root trait space** |  |  |  |  |  |  |  |  |  |
| --- | --- | --- | --- | --- | --- | --- | --- | --- | --- |
|  | **Dissimilarity** |  |  | **Overlap** |  |  | **Not shared** |  |  |
|  | **PC** | **PNC** | **W** | **PC** | **PNC** | **W** | **PC** | **PNC** | **W** |
| **A** | 0.34 (0.05) | 0.38 (0.04) | 0.48 (0.04) | 0.83 (0.06) | 0.83 (0.08) | 0.86 (0.07) | 0.17 (0.06) | 0.17 (0.08) | 0.14 (0.07) |
| **PC** |  | 0.28 (0.05) | 0.33 (0.04) |  | 0.88 (0.08) | 0.88 (0.05) |  | 0.11 (0.07) | 0.12 (0.05) |
| **PNC** |  |  | 0.33 (0.04) |  |  | 0.90 (0.04) |  |  | 0.10 (0.04) |

### Linking clonal traits to aboveground and fine-root trait combinations

Clonal and bud bank traits were significantly associated with aboveground trait combinations (**Fig. 1a-d**, p always < 0.05, full model statistics in **Table S3**) but the patterns differed among clonal traits and the relationships had a generally low explanatory power (R^2^ = 4-22%). For all data pooled, total bud-bank size and lateral spread were associated with the greatest plant height, seed mass and leaf area (hot spot in **Fig. 1a**,**c**). Multiplication rate increased towards lower values of plant height, seed mass and leaf area and towards greater values of specific leaf area and leaf nitrogen content (hot spot in **Fig. 1b**), but this pattern was overall very weak. Persistence of clonal connection increased with specific stem density (hot spots in **Fig. 1a**,**b**,**d**). Belowground (**Fig. 2a-d, Table S3**), only two significant relationships were found: lateral spread increased with the N content in fine roots (hot spot in **Fig. 2c**, R^2^ = 5%, p = 0.004), and persistence of clonal connection increased with root tissue density (hot spot in **Fig. 2d**, R^2^ = 0.10, p = 0.002). However, the patterns of how clonal related traits varied within the aboveground and fine-root trait space sometimes differed between functional groups (**Appendix S2**; **Fig. S12-S19, Table S3**). In addition, we found a significant but overall weak correlation between clonal-trait combinations (i.e., PCA run on clonality-related traits) and the aboveground or fine-root trait space of clonal plants. In detail, the correlation between the clonality-trait space and the aboveground trait space was 0.26 (Procrustes Sum of Squares = 0.93, p = 0.001), while that of clonality-trait space and the fine-root trait space was 0.20 (Procrustes Sum of Squares = 0.96, p = 0.003).

**Fig. 1.**
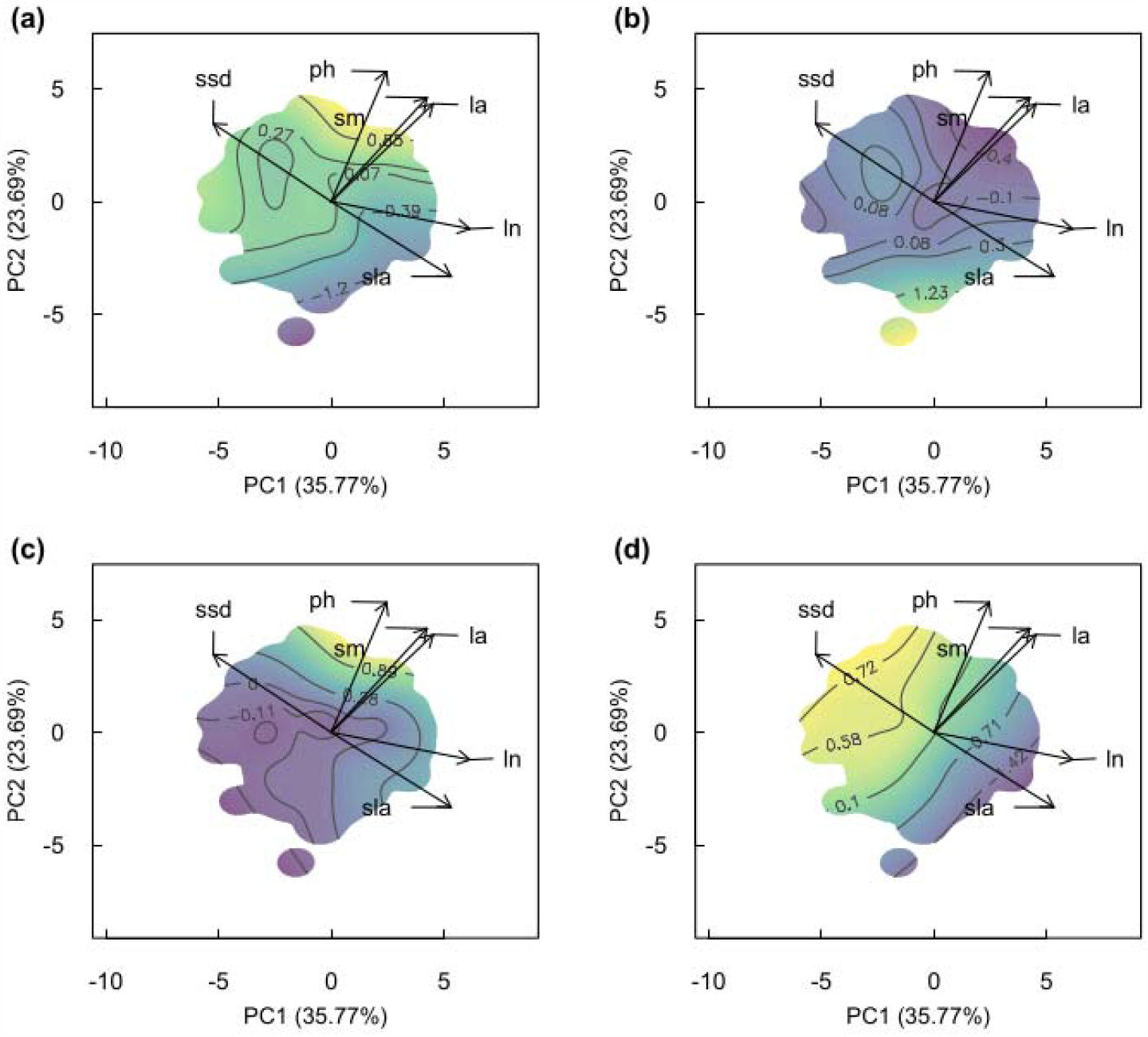
Mapping clonality-related traits in the aboveground trait space. **(a)** Total bud-bank size; **(b)** Multiplication rate; **(c)** Lateral spread; **(d)** Persistence of clonal connection. High values of the trait being mapped are shown in yellow. See Methods for details on the methodology. ssd = specific stem density, ph = plant height, sm = seed mass, la = leaf area, ln = leaf nitrogen content (mass basis), sla = specific leaf area. Details for the considered traits are shown in **Table S1**. Grey lines within the map correspond to quantiles of GAM predictions. Note that the response variable was scaled before the analysis. Full GAM statistics are reported in **Table S3**.

**Fig. 2.**
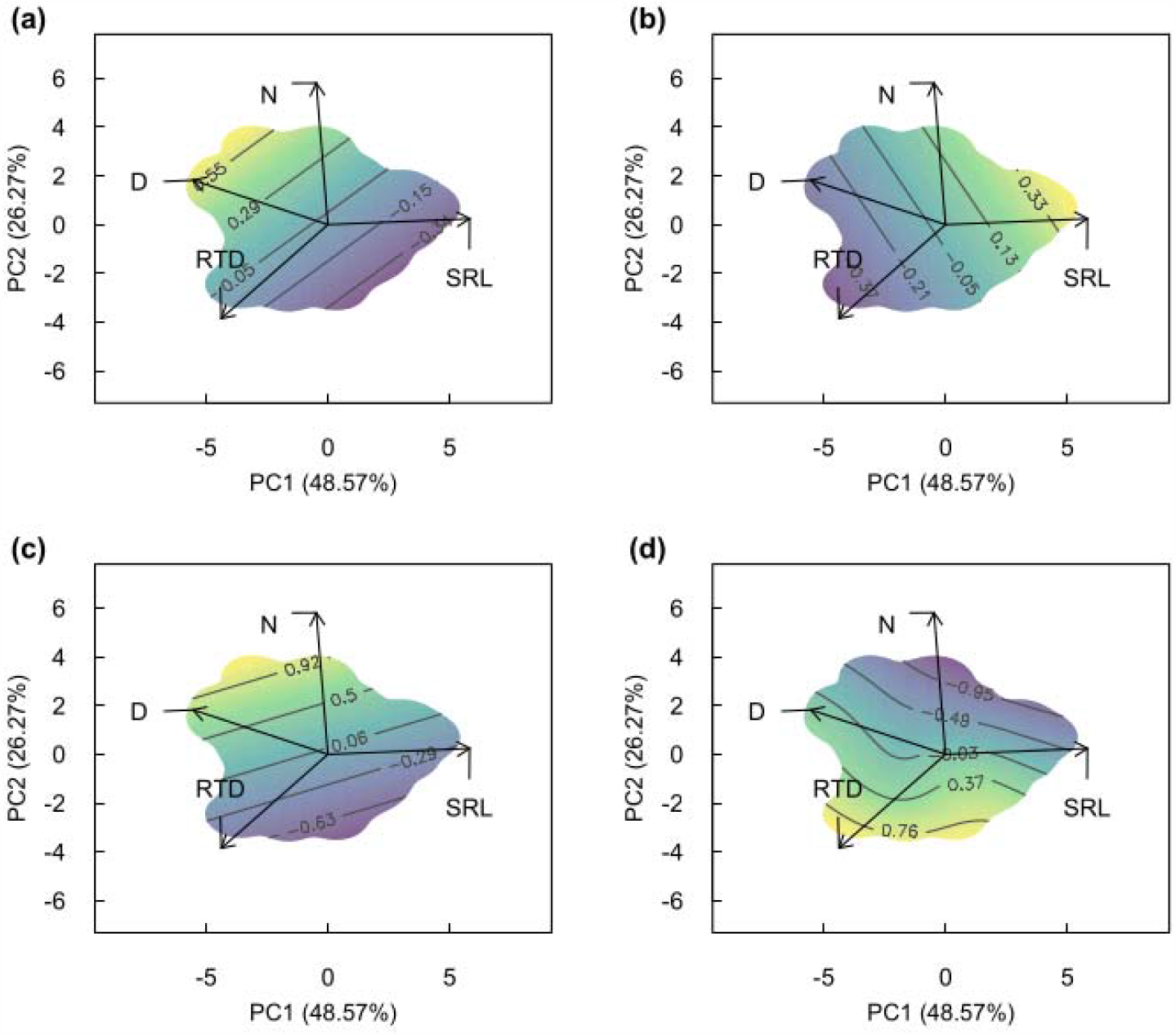
Mapping clonality-related traits in the fine-root trait space. **(a)** Total bud-bank size; **(b)** Multiplication rate; **(c)** Lateral spread; **(d)** Persistence of clonal connection. High values of the trait being mapped are shown in yellow. See Methods for details on the methodology. D = mean root diameter, N = root nitrogen content (mass basis), SRL = specific root length, RTD = root tissue density. Details for the considered traits are shown in **Table S1**. Grey lines within the map correspond to quantiles of GAM predictions. Note that the response variable was scaled before the analysis. Full GAM statistics are reported in **Table S3**.

## Discussion

We provide the most comprehensive analysis to date of how clonal and bud-bank traits integrate in the aboveground (Díaz et al. 2016) and fine-root (Carmona et al. 2021) trait spaces. We found that clonal herbs cannot be distinguished from non-clonal herbs only based on their aboveground or fine-root traits. Furthermore, we spotlight a weak coordination between clonal traits and both aboveground and fine-root trait combinations in clonal perennial herbs. Altogether, our findings suggest that clonality provides perennial herbs with important additional functions not mirrored by the other trait dimensions like horizontal space acquisition, space holding ability, vegetative regeneration and clonal multiplication. Thus, clonality provides independent axes of specialization in perennial herbs.

### Aboveground and fine-root trait dimensions do not distinguish clonal from non-clonal herbs

Perennial clonal and non-clonal herbs were indistinguishable by using the available aboveground and fine-root trait combinations. Being clonal, plants are not constrained in their acquisitive traits, size (measured as plant height and leaf size), or investments into generative reproduction (expressed as seed mass). They differ in characters that were so far poorly represented in trait-based ecology, for example, size in horizontal dimension, number of connected rooting units, storage organs size, regeneration capacity with bud bank. Thus, clonality might provide perennial herbs with an additional way to persist, exploit resources in space and deal with disturbance. Clonal and aboveground and fine-root traits might be subjected to different selective pressures, as already suggested for example for leaf economics and floral traits (Murren 2002; Zhang et al. 2017). For example, clonal plants can cope with spatial and temporal resource variation by iterating the basic units from which they are constructed (Hutchings & Bradbury 1986). Such iteration of the basic units would allow clonal plants to spatially and temporally change their positioning along (micro-)environmental gradients to reach the spots where their trait combinations maximize performance, or by diversifying trait combinations through their high degree of phenotypic plasticity (relative to non-clonal plants; Herben & Klimešová 2020) in the new reached location. These strategies might not be mutually exclusive, and they could interact to define a variety of possible trait combinations energetically supported by the resource sharing among ramets on the move (Jónsdóttir & Watson 1997). Besides, the acquisition of the clonal reproduction is quite flexible, being it a character that can be easily gained or loss (Herben & Klimešová 2020). If aboveground or fine-root traits change at a slower rate through evolutionary times compared to clonal reproduction (and its associated traits), this might further explain why clonal and not-clonal herbaceous plants do not differ in the amount of possible trait combinations. A similar explanation was provided to explain, for instance, the independence of vegetative and floral traits (Opedal 2019). Overall, by comparing clonal and non-clonal herbs allowed us to identify that being clonal does not constraint aboveground or fine-root functional differentiation in perennial herbs. This likely depends on the interaction between the ecological and evolutionary reasons outlined above, and contribute to explain the large independence between clonal traits and aboveground and fine-root trait combinations (Discussed below).

### Clonal and bud bank traits are largely decoupled from aboveground and fine-root trait spectra

When focusing on perennial clonal herbs only, we found an overall weak coordination between clonality-related traits and the aboveground and fine-root trait spectra. This hints that clonal and bud bank traits might represent an independent axis of functional specialization, generalizing previous findings based on more limited species sets (Herben et al. 2012, 2016, 2018; Klimešová et al. 2016), and supporting our previous considerations. In addition, the observed patterns sometimes varied between the functional groups of clonal forbs and graminoids, suggesting that clonality related traits are equally successful in contrasting ecological contexts (broadly reflected by the known differences in ecological requirements between forbs and graminoids; Zhang et al. 2023). This pattern also suggests that some of the observed relationships on pooled data might be pertinent only to some clonal species belonging either to forbs or graminoids and might depend on the clonal trait under examination. For example, some hotspots might appear under specific environmental regimes (e.g., highly disturbed habitats), or reflect an allometric scaling with size-related traits. In other words, some very specific associations between aboveground and fine-root trait combinations with clonal traits might not be always linked to being clonal *per se*.

Aboveground, despite being characterized by a low explanatory power, lateral spread and bud bank size were positively related to plant height, seed mass and leaf area which overall mirror the size-related dimension of the global spectrum of plant form and function. This means that species with higher lateral spread and larger bud bank are large plants with great organs’ size. This might reflect an allometric scaling (Klimešová et al., 2016). However, regarding lateral spread, this result contrasts with previous findings in which this trait (i) was positively related to aboveground traits mirroring the resource economics dimension (e.g., specific leaf area, Klimešová et al., 2016), and (ii) was negatively related to plant size (i.e., plant height and seed mass; Ladouceur et al., 2019; Lubbe et al., 2021; but see Klimešová et al., 2016). The larger array of strategies, and thus of trait ranges, included in our study can partly explain these discrepancies compared to smaller cross-species studies often carried out at local scales. Besides, we want to stress that when combining traits within trait spaces, some expected correlations might disappear due to the shift in coordinates of species from a bivariate to a multivariate space. For example, a given correlation between two traits found in a previous study might be due to the covariance of these two traits with a third player (Puglielli et al. 2021).

The most consistent relationship we found both on pooled data, for each functional group, and at any given plant and organ size, was the positive scaling between persistence of clonal connection and specific stem density (sometimes approximated by leaf dry matter content in herbaceous plants, Díaz et al. 2022), associated to smaller specific leaf area and leaf nitrogen content. This pattern implies that persistence of clonal connection is an economic trait for clonal plants, as slow growing plants with longer leaf lifespans have usually long-lived connections between ramets (Klimešová et al. 2016). Additionally, the positive scaling between persistence of clonal connection and specific stem density might reflect the fact that more persistent connections are ensured by stem-derived organs with higher mechanical safety and resistance against pathogens, herbivores or physical damage (e.g., hypogeogenous rhizomes).

Belowground, lateral spread and persistence of clonal connection slightly varied along the spectrum defined by fine-root root nitrogen content and tissue density (Carmona et al. 2021). On the one hand, lateral spread was positively associated with high nitrogen content, probably implying that horizontal clonal spreading is partially related to acquisitive fine-root strategies. However, we call for cautiousness when interpreting this pattern, as it was very weak and strongly dependent on the considered functional group (possibly owing to the presence of Leguminosae). On the other hand, and again, persistence of clonal connection was the only trait showing a strong pattern and it was positively associated with root tissue density, confirming that plants living in nutrient and water limited environments not only have conservative fine-root strategies, but also persistent clonal organs (e.g., rhizomes) ensuring long-term resource sharing and storage (Jónsdóttir & Watson 1997). Overall, the persistence of clonal connection is the only clonal trait mirroring a trade-off between fast and slow return of investment both above- (i.e., stem specific density) and belowground (i.e., root tissue density), as already discussed.

### Towards a clonal-trait space

The weak correlation between the clonality-trait space and both the aboveground and fine-root trait spaces means that the clonal trait space is only partially determined by leaf and root economics and by organs’ size, as already discussed. This indicates that the investment in clonal growth only partially depends on resource acquisition strategies and plant size. As such, species positioning in the clonal trait space is largely uncorrelated with species positioning in the aboveground or fine-root trait space, meaning that the clonal trait space is largely independent from the other spaces. Indeed, the investment in the maintenance of clonal connections implies that, when ramets die back, the resources they have accumulated are not lost, but can be translocated for subsequent use elsewhere in a clone. Thus, accumulated resources might be redistributed from older to younger parts of the clone without implications for resource acquisition strategies (Hutchings & Bradbury 1986). On the other hand, clonal plants are usually characterized by a lower investment in seed production (Herben et al. 2015) and rooting depth (Šmilauerová & Šmilauer 2007; Klimešová & Herben 2023) with respect to non-clonal plants, thus limiting their ability to quickly colonize newly created habitats in early succession and to inhabit dry habitats (Latzel et al. 2011). Besides, clonal plants might always favor horizontal over vertical growth, decoupling the horizontal from the vertical size dimension (Klimešová et al. 2018).

From previous considerations it becomes evident that the functions captured by clonal trait dimensions does not mirror the same functions ascribed to above- and fine-root axes of specialization. We therefore explored the potential clonal specialization axes and identified two leading dimensions summarizing clonal and bud bank trait combinations in clonal perennial herbs (**Fig. 3**). The first dimension (explaining 38.9% of the total variance) reflects a positive scaling between bud bank size and persistence of clonal connection, and it can be interpreted as a “*persistence”* axis. This axis might reflect a spectrum of strategies from splitting clones (splitter-integrator continuum *sensu* Jonsdottir & Watson 1997) with small bud bank towards integrated clones with large bud bank typical for unproductive habitats with intermediate disturbance frequency (Bellingham & Sparrow 2000; Herben et al. 2018). The second dimension (explaining 32.0% of the total variance) is defined by the positive scaling between multiplication rate and lateral spread, and it can be interpreted as a “*clonal multiplication*” axis. This axis mirrors the capacity to occupy new space by growing clonally and might represent a trade-off between competitive ability *vs*. stress tolerance. Consequently, it is related to resource availability which might be higher where we find higher multiplication rates and lateral spread, and lower at the other hand of the axis. However, lateral spread has other functions, such as the fine-scale exploration of new habitats when resources are heterogeneously distributed in space and time (Klimešová□□ et al. 2018). Overall, our results suggest that clonal trait combinations might provide alternative ways to endure stress through different mechanisms than those provided by acquisitive traits (both above and belowground) and to endure disturbance through other regenerative traits than seeds. The advantages that such alternative strategies might bring under multiple ecological contexts in comparison to aboveground and fine-root trait combinations need further investigation.

**Figure 3.**
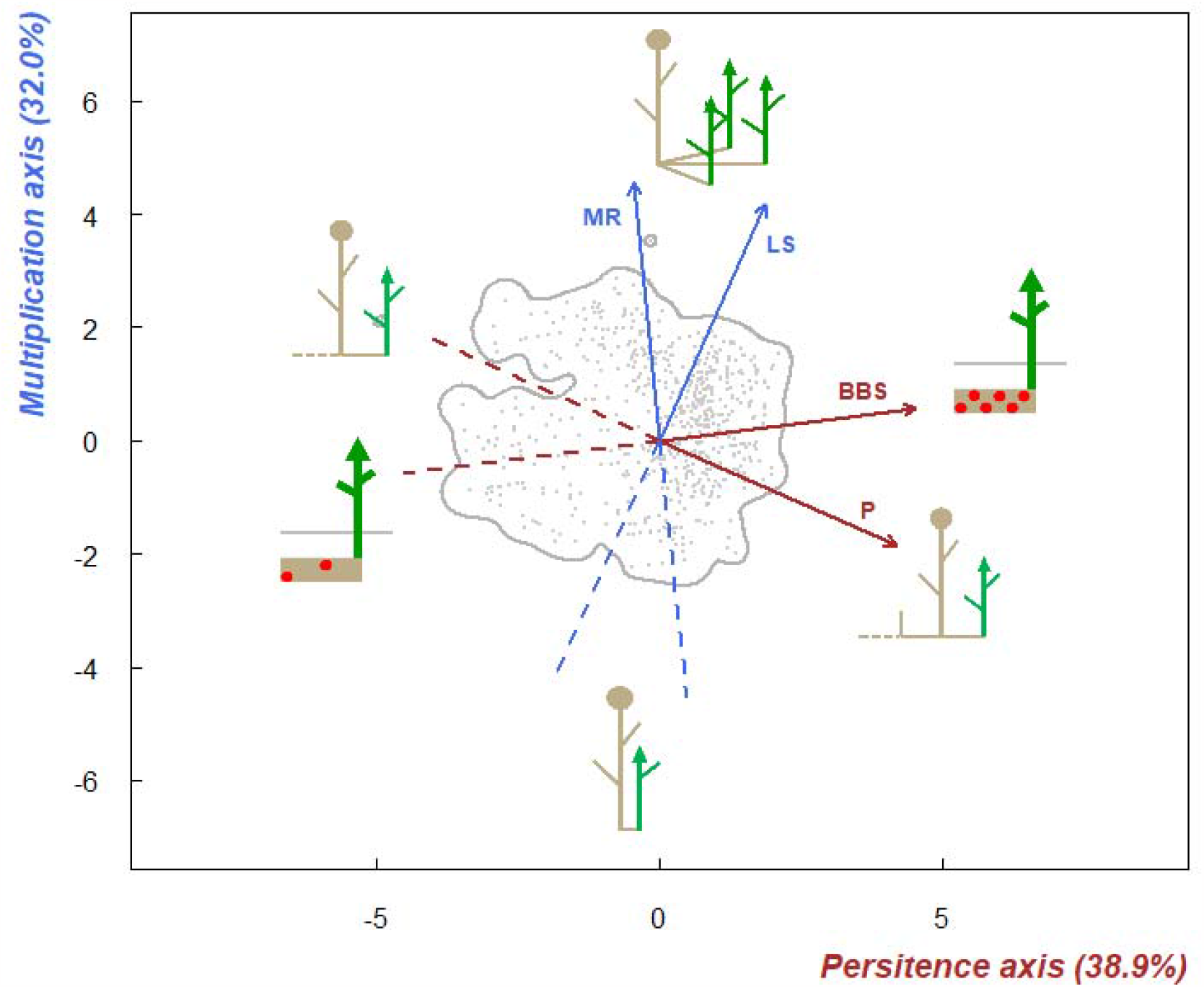
Towards a clonal trait space. The trait space defined by a PCA on clonal trait data for 1181 clonal species with complete trait information. The first component is interpreted as a ‘*persistence axis*’ running from splitting clones with short persistence of clonal connections (P) and small bud bank size (BBS) to integrated clones with longer P and larger BBS. The second component is interpreted as a ‘*multiplication axis*’, a spectrum from low to high rates of clonal multiplication (MR) and lateral spread (LS). The percentage of variance in the data that is explained by each axis is shown. The grey line represents the 0.99 quantile of the multivariate kernel density function. Drawings from Klimešová et al. (2016).

## Conclusions

We provide the first evidence that at a large taxonomic scale clonal traits add independent trait dimensions to the plant spectrum of forms and functions. However, despite the eco-evolutionary relevance of clonal and bud bank traits and the availability of standard protocols for their sampling (see Klimešová et al. 2019), there is still a huge data gap for species outside Europe (Klimešová et al. 2017). This potential geographical and functional limitation of our study - which is indeed limited to perennial herbs from Central Europe - can however turn into an advantage as our results can be considered spurious of many potential confounding effects typical of large cross-species studies (e.g., species distribution across disparate biomes, broad latitudinal gradients, many functional types). Finally, as a natural next step towards defining the clonal trait space, we call for the need of a global assessment of the patterns highlighted in this study. For this purpose, ecologists need to go belowground and measure clonal and bud bank traits together with other functional traits, at least across those biomes where such traits are recognized to be fundamental for plant strategies and therefore for ecosystem functioning (e.g., grassy and shrubby biomes; Ottaviani et al. 2020).

## Acknowledgements

J.K. acknowledges support by the Praemium Academiae award from the Czech Academy of Sciences of the Czech Republic.

J.L.T. was supported by the LIFE MODERn (NEC) project (LIFE20 GIE/IT/000091).

G.P. was supported by the grants IJC2020-043331-I funded by MCIN/AEI/10.13039/501100011033, and the grant PID2021-122214NA-I00 funded by MCIN/AEI/ 10.13039/501100011033 and by FEDER “ESF Investing in your future”.

## Author contributions

S.C. and G.P. conceived the idea. S.C., J.K. and GP conceptualized the study. G.P and J.L.T. performed statistical analyses. S.C., and G.P. led the writing of the paper with inputs from J.K. and J.L.T. All authors read and approved the final manuscript.

