## Supplementary material for "Clonality-Related Traits Add Independent Specialization Axes to Herbs’ Trait Strategies"

**Supplementary information**

**Table S1. List of traits used, their abbreviation (in parenthesis), description and related function.**

| Trait | Unit | Definition | Function |
| --- | --- | --- | --- |
| Plant height (PH) | m | Maximum height of a whole plant | Ability to pre-empt light resources and disperse diaspores |
| Specific stem density (SSD) | mg mm^−3^ | Ratio of the mass of the stem after drying to its volume assessed without drying | Trade-off between growth potential and mortality risk from biomechanical or hydraulic failure |
| Seed mass (SM) | mg | Mass of a seed assessed after drying | Trade-off between seedling survival versus colonization ability in space and time |
| Leaf size (LA) | mm^2^ | Area of a leaf in the one-sided projection | Important consequences for leaf energy and water balance |
| Specific leaf area (SLA) | mm^2^ mg^-1^ | Ratio of the area of a leaf to its dry mass | Trade-off between carbon gain and longevity |
| Leaf nitrogen content (LN) | mg g^-1^ | Ratio of the quantity of nitrogen in the leaf per respective unit dry mass | Trade-off between the benefits of photosynthetic potential and the costs of acquiring nitrogen and suffering herbivory |
| Specific root length (SRL) | m g^-1^ | Ratio of the length of fine roots to the corresponding fine root mass assessed after drying | Important for resource acquisition.  It reflects the potential extent of soil exploration per unit cost (in terms of root biomass allocation) |
| Mean root diameter (D) | mm | Diameter of a root | Trade-off between resource acquisition and stress tolerance (or mycorrhizal colonization) |
| Root nitrogen content (N) | mg g^-1^ | Ratio of the quantity of nitrogen in the root to the corresponding root mass assessed after drying | Informative on plant metabolic rate |
| Root tissue density (RTD) | g cm^-3^ | Ratio of root mass after drying to the volume assessed without drying | Trade-off between growth rate and longevity |
| Total bud-bank size (BBS) | - | Total number of buds stored in bud bearing organs per rooting unit | Recovery after damage; Space occupancy; On-spot persistence |
| Multiplication rate (MR) | - | Number of offspring rooting units produced by a parental rooting unit per year | Space occupancy; On-spot persistence; Competitive ability |
| Lateral spread (LS) | m year^-1^ | Distance between offspring rooting units and parental rooting unit | Space occupancy; Resource foraging; Competitive ability |
| Persistence of clonal connection (P) | year | Period of connection between offspring rooting unit and parental rooting unit | On-spot persistence; Competitive ability |

**Appendix S1. Building aboveground and fine-root trait spaces and testing the effect of trait data imputation**

We used Principal Component Analysis (PCA) to visualize the trait spaces defined by the combinations of the aboveground (**Fig. S1**) and fine-root traits (**Fig. S2**). Before running PCA we imputed missing data using the ‘missForest’ R package (Stekhoven & Bühlmann, 2012). The function uses trait-trait relationships combined with species phylogenetic relatedness (as proposed by Penone et al., 2014) to impute missing data. Phylogenetic data were retrieved for all the species in our above- and belowground dataset using the V.PhyloMaker R package (Jin & Qian, 2019). V.PhyloMaker uses a mega-tree that combines the phylogenies developed by Zanne et al. (2014) and Smith & Brown (2018). This imputation procedure has been already proved to yield solid results with very low error rates (e.g., 1.4%, Carmona et al., 2021). Overall, the imputation procedure did not alter trait-trait relationships nor the definition of the above- and belowground trait spaces (**Appendix S1; Fig. S3-S4: Table S2**). The imputation procedure returned datasets with a completeness of 100% for all the considered traits (2655 and 430 observations for the above- and belowground trait space, respectively).

In most of the cases, the imputation procedure did not alter the trait-trait relationships for aboveground (**Fig. S3a-q**) or fine-root traits (**Fig. S4a-f**). The only exceptions were the relationship SSD-SM for the aboveground compartment, and the relationship SRL-N and N-RTD that changed slope in the imputed and not-imputed fine-root dataset (**Fig. S4c**). This can be explained by an overall weak relationship between the considered traits. SSD and SM are independent in the aboveground trait space. Similarly, within the fine-root trait space, N is independent of SRL, and negatively related to RTD, which in turn tends to be positively related to D, is the latter being independent of N. These patterns of covariation might complicate the imputation of any of these traits. To test the effect of this problematic imputation cases on the defined trait spaces, we employed the following procedure: (i) we removed one target trait at a time from the considered trait space; and (ii) we quantified the correlation between the observed and each reduced trait space using Procrustes analysis. This analysis rotates one configuration (the observed fine-root trait space) to maximum similarity with a target one (any of the reduced fine-root trait space) and tests the non-randomness of the similarity via permutation (999 randomizations) based on Monte-Carlo simulations. Since for fine-root traits we found imputation issues for relationships involving three of the four traits used to define the trait space, we applied the above-described procedure by removing all the traits once at a time. The results returned a mean correlation coefficient of 0.925±0.02 between reduced and full trait spaces (**Table S2**), indicating that the imputation issues are completely resorbed by trait covariation patterns within the aboveground and fine-root trait space.

**Fig. S1. Trait space defined by aboveground traits for all the species included in the dataset.** The dataset spanned annual (n = 577), perennial clonal (1323), perennial non-clonal (603), and woody (152) plants. Colored areas refer to the probability of finding some trait combinations within the trait space (red area = high probability). Contour lines show the 0.99, 0.50 and 0.25 quantile of the multivariate probability distribution of trait combinations (see Methods for further details). Trait abbreviations are reported in **Table S1**.

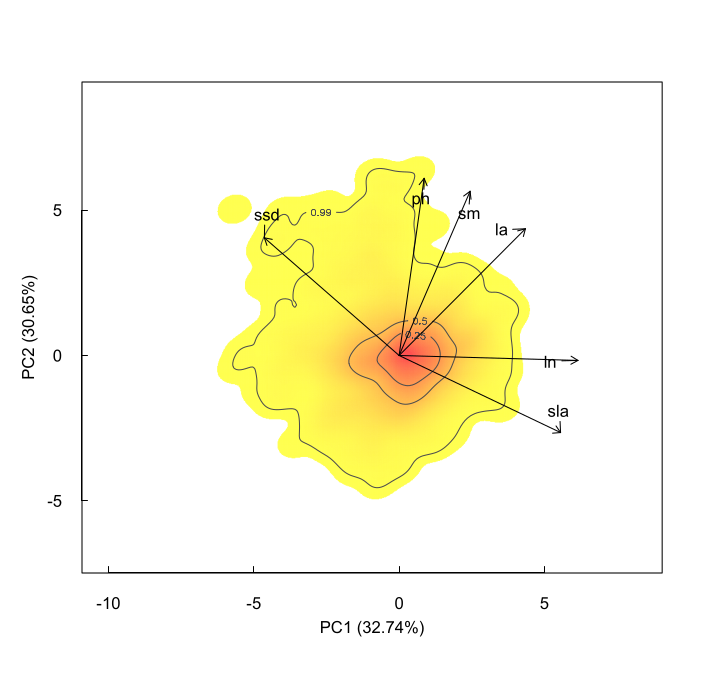

**Fig. S2. Trait space defined by fine-root traits for all the species included in the dataset.** The dataset spanned annual (n = 90), perennial clonal (195), perennial non-clonal (79), and woody (66) plants. Colored areas refer to the probability of finding some trait combinations within the trait space (red area = high probability). Contour lines show the 0.99, 0.50 and 0.25 quantile of the multivariate probability distribution of trait combinations (see Methods for further details). Trait abbreviations are reported in **Table S1**.

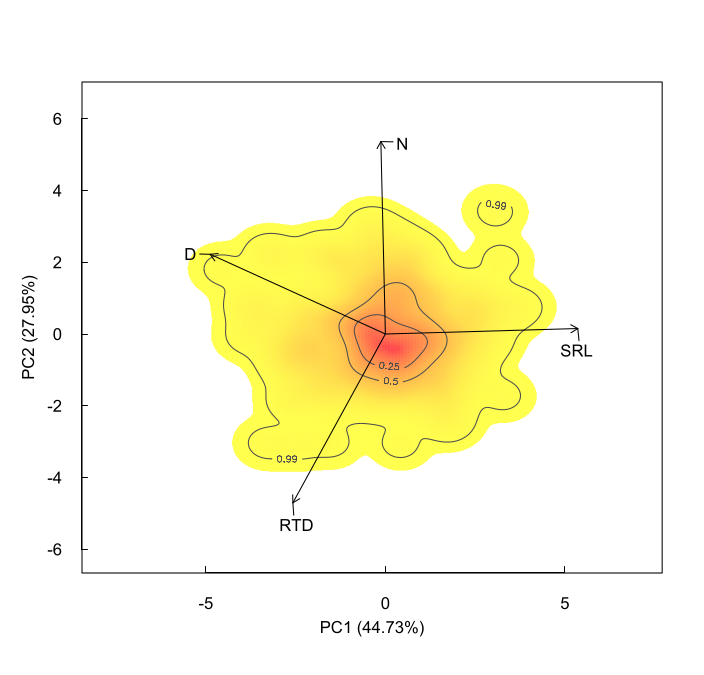

**Fig. S3. Testing the effect of imputation on aboveground trait-trait relationships.** The test was carried out by comparing changes in the trait-trait relationships between the imputed (blue symbols) and not-imputed dataset (red symbols). The fitted lines represent Standardized Major Regressions carried out using the ‘smatr’ R package (Warton et al. 2012).

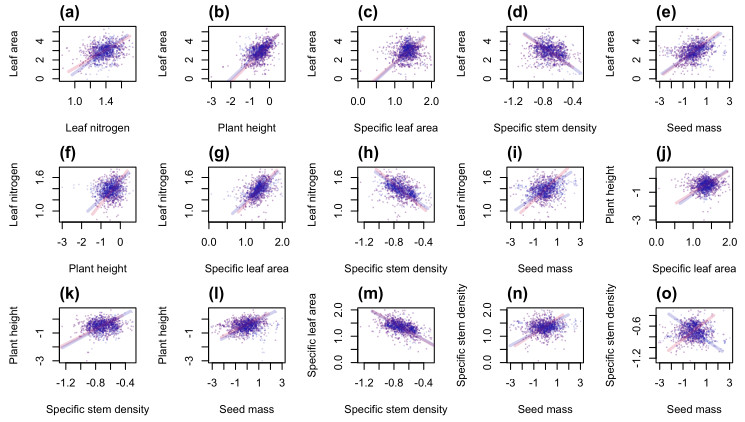

**Fig. S4. Testing the effect of imputation on belowground trait-trait relationships.** The test was carried out by comparing changes in the trait-trait relationships between the imputed (blue symbols) and not-imputed dataset (red symbols). The fitted lines represent Standardized Major Regressions carried out using the ‘smatr’ R package (Warton et al. 2012).

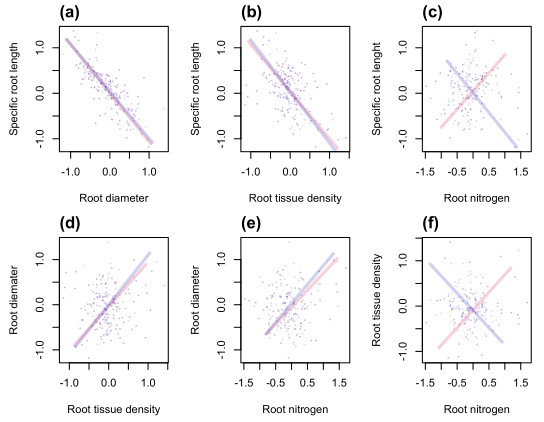

**Table S2. Testing the effect of the problematic imputation cases on the definition of aboveground and fine-root trait spaces.** This test was executed by removing one trait with problematic imputation results at a time from the considered trait space; and by quantifying the correlation between the observed and each reduced trait space using Procrustes analysis (See **Appendix 1** for further details). High values of the correlation coefficients indicate very good correspondence between trait spaces.

| **Observed aboveground trait space vs.** | **Space obtained removing:** | **Correlation** | **p-value** |
| --- | --- | --- | --- |
|  | SSD | 0.94 | 0.001 |
|  | SM | 0.92 | 0.001 |
| **Observed belowground trait space vs.** | **Space obtained removing:** | **Correlation** | **p-value** |
|  | N | 0.96 | 0.001 |
|  | SRL | 0.87 | 0.001 |
|  | D | 0.95 | 0.001 |
|  | RTD | 0.91 | 0.001 |

**Fig. S5.** Distribution of annual (n = 577), perennial clonal (1323), perennial non-clonal (603), and woody (152) plants in the aboveground trait space (grey line, **Fig. S1**). Pairwise comparisons of the TPD overlap-based dissimilarity are shown in **Table 1** of the main text. The dashed line represents the trait space defined across growth forms. According to PERMANOVA analysis carried out using either PC1 or PC2 as the response variable and growth form as the grouping variable, the latter explained 6% of traits variation along PC1 and 31% along PC2. For trait abbreviations see **Table S1**.

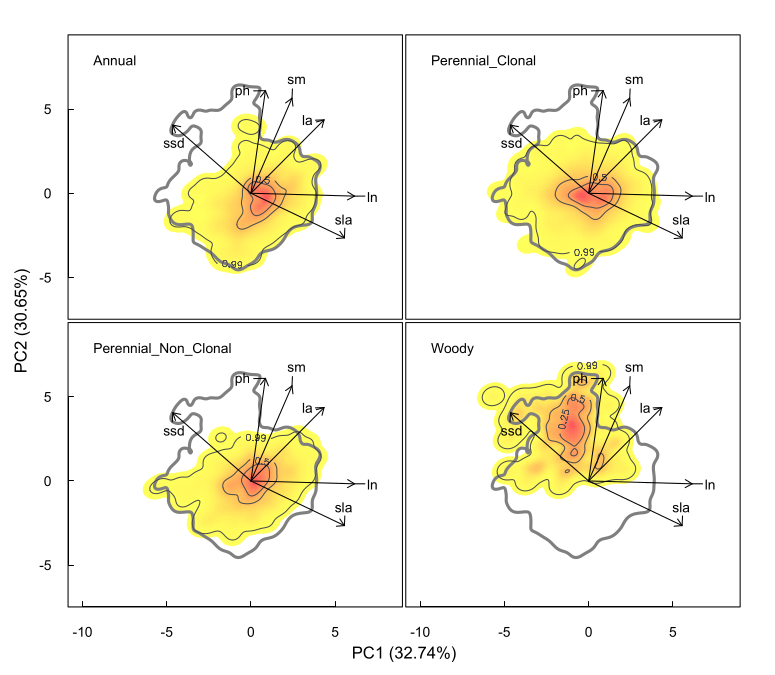

**Fig. S6.** Distribution of annual (n = 90), perennial clonal (195), perennial non-clonal (79), and woody (66) plants in the fine-root trait space (grey line, **Fig. S1**). Pairwise comparisons of the TPD overlap-based dissimilarity are shown in **Table 1** of the main text. The dashed line represents the trait space defined across growth forms. According to PERMANOVA analysis carried out using either PC1 or PC2 as the response variable and growth form as the grouping variable, the latter explained 7.5% of traits variation along PC1 and 3.2% along PC2. For trait abbreviations see **Table S1**.

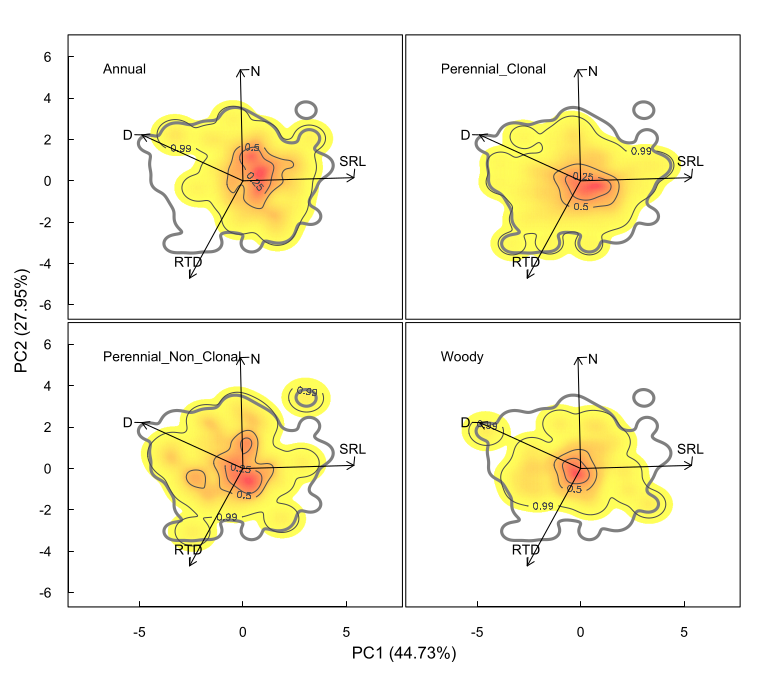

**Fig. S7. Trait space defined by aboveground traits for the clonal species in the dataset.** Colored areas refer to the probability of finding some trait combinations within the trait space (red area = high probability). Contour lines show the 0.99, 0.50 and 0.25 quantile of the multivariate probability distribution of trait combinations (see Methods for further details). Trait abbreviations are reported in **Table S1**.

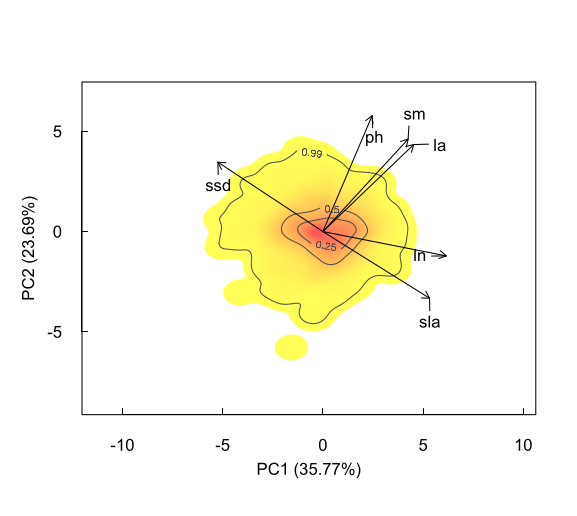

**Fig. S8. Trait space defined by fine-root traits for the clonal species in the dataset.** Colored areas refer to the probability of finding some trait combinations within the trait space (red area = high probability). Contour lines show the 0.99, 0.50 and 0.25 quantile of the multivariate probability distribution of trait combinations (see Methods for further details). Trait abbreviations are reported in **Table S1**.

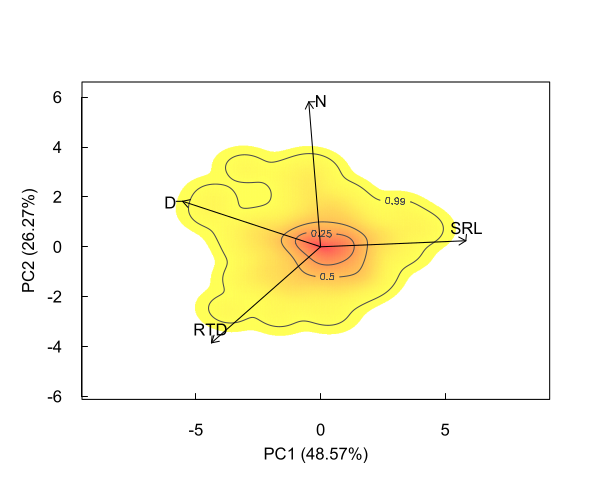

**Appendix S2. Trait space occupancy and mapping of clonal traits in the aboveground and fine-root trait spaces of clonal forbs and graminoids**

After having defined the trait spaces for clonal plants (see **Linking clonal traits to aboveground and fine-root trait combinations** section in the main text), we analyzed how clonal plants belonging to the two functional groups in our dataset, forbs and graminoids, differed in their positioning within the trait spaces (**Fig. S9-S11**). We then analyzed: (i) the extent of the shared portion of the trait space between the two groups (i.e., overlap-based dissimilarity); (ii) the main functional diversity indexes, namely functional richness, evenness, and divergence, following Carmona et al. (2016) (see also Mason et al. 2005). Functional richness quantifies the trait space area occupied by each group. Functional evenness reflects the evenness in the abundance of trait combinations in each group. Functional divergence reflects how far some trait combinations are from the center of gravity of the space per each group. Overlap-based dissimilarity and functional diversity indexes were calculated using kernel-density-based functions included in the TPD R package (Carmona et al., 2019). However, considering that the two considered groups differ in their sample size (919 vs. 338, and 110 vs. 85 forbs and graminoids in the aboveground and fine-root dataset, respectively), we assessed the overlap-based dissimilarity and the functional diversity indexes by using an equally sized random sample of the species in each group (n = 50 for above- and belowground traits) across 999 iterations.

Clonal forbs and graminoids differed in their positioning within aboveground and fine-root trait spaces. Aboveground, TPD-based dissimilarity was 0.54 ± 0.06 with a trait space overlap of 0.83 ± 0.08, and a not-shared portion of the of 0.17 ± 0.07. Belowground, TPD-based dissimilarity between forbs and graminoids was 0.36 ± 0.05, with an overlap and a not-shared portion of the trait space of 0.85 ± 0.07 and 0.15 ± 0.07, respectively. Forbs displayed a greater functional richness, functional evenness, and functional divergence both above- and belowground compared to graminoids (**Fig. S11a-f**, p always < 0.001, n = 999, t-test). Altogether, these results indicate that, despite large overlap, the density hot- and cold-spots of trait combinations tend to differ between forbs and graminoids, and they differently occupy the trait spaces.

Forbs and graminoids with the greatest values of total bud bank size displayed greater values of plant height, seed mass and leaf area (hot spot in Fig. S6a, see Data analysis section in the main text) and a tendency towards higher specific stem density compared to the species with the lowest total bud bank size (hot spot in **Fig. S11**). However, for graminoids the pattern was less evident than for forbs.. Multiplication rate was associated with greater values of specific leaf area and leaf nitrogen in forbs, associated to the lowest values of plant height, seed mass and leaf area (hot spot in **Fig. S12**) but the pattern was not significant for graminoids (hot spot in **Fig. S12**). Lateral spread increased with plant height, leaf area and seed mass in both forbs and graminoids (hot spots in **Fig. S13**). Persistence increased with SSD in both functional groups (**Fig. S6g,h**). Belowground, most of the relationships were not significant within each functional group **(Fig. S15-18**). Increasing lateral spread with increasing RTD and D for graminoids was the only significant relationship (**Fig. S17f**).

**Fig. S9. Distribution of clonal forbs and graminoids in the aboveground trait space.** Colored areas refer to the probability of finding some trait combinations within the trait space (red area = high probability). Contour lines show the 0.99, 0.50 and 0.25 quantile of the multivariate probability distribution of trait combinations (see Methods for further details). Trait abbreviations are reported in **Table S1**.

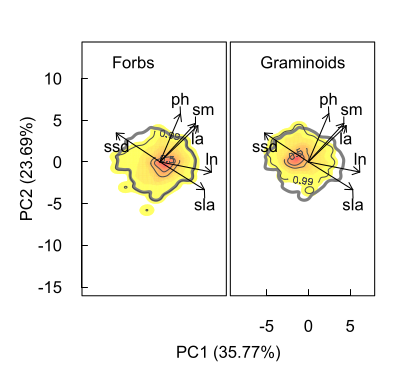

**Fig. S10. Distribution of clonal forbs and graminoids in the fine-root trait space.** Colored areas refer to the probability of finding some trait combinations within the trait space (red area = high probability). Contour lines show the 0.99, 0.50 and 0.25 quantile of the multivariate probability distribution of trait combinations (see Methods for further details). Trait abbreviations are reported in **Table S1**.

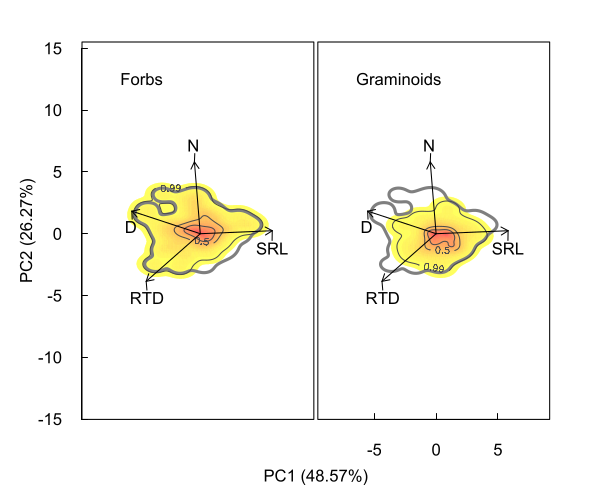

**Fig. S11. Comparing functional diversity between clonal forbs and graminoids.** Functional diversity indexes for the **(a-c)** above- and **(d-f)** belowground trait space. Functional richness summarizes the area/volume occupied by a trait space, functional evenness represents the extent by which trait combinations are evenly distributed within a trait space, and functional divergence, quantifies the density of points that are far from the trait space’s center of gravity. Each grey point represents the average value of 1000 iterations at the same sample size for the two groups (see **Appendix S2**).

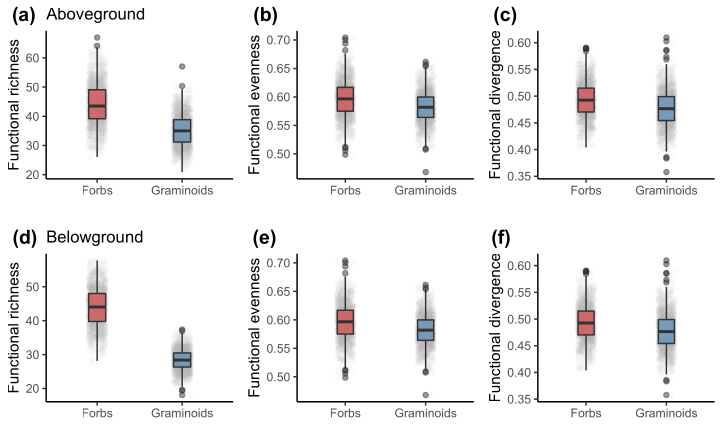

**Fig. S12. Mapping total bud-bank size in the aboveground trait space for forbs (left column) and graminoids (right column).** Details on the mapping can be found in **Appendix S2.** Trait abbreviations as in **Table S1**.

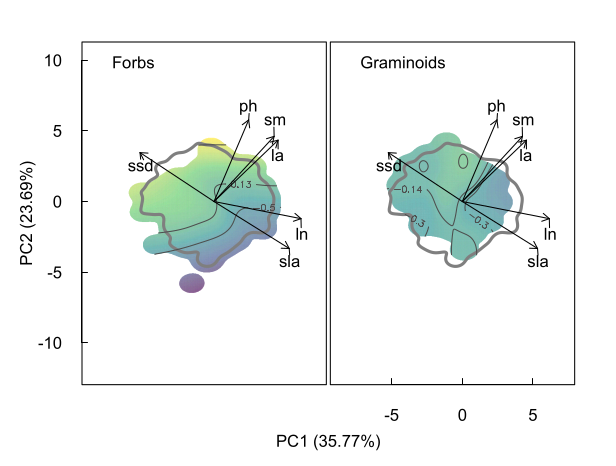

**Fig. S13. Mapping multiplication rate in the aboveground trait space for forbs (left column) and graminoids (right column).** Details on the mapping can be found in **Appendix S2.** Trait abbreviations as in **Table S1**.

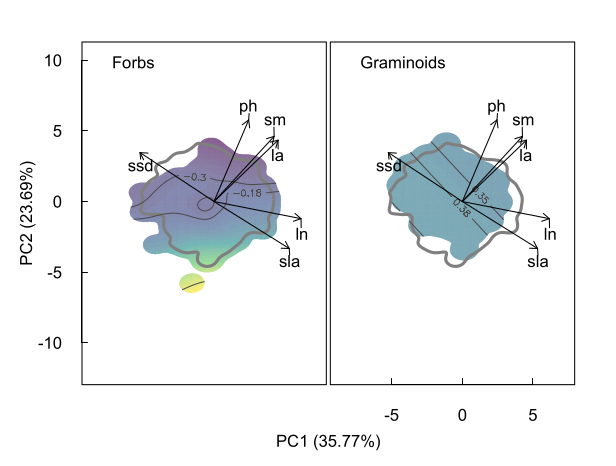

**Fig. S14. Mapping lateral spread in the aboveground trait space for forbs (left column) and graminoids (right column).** Details on the mapping can be found in **Appendix S2.** Trait abbreviations as in **Table S1**.

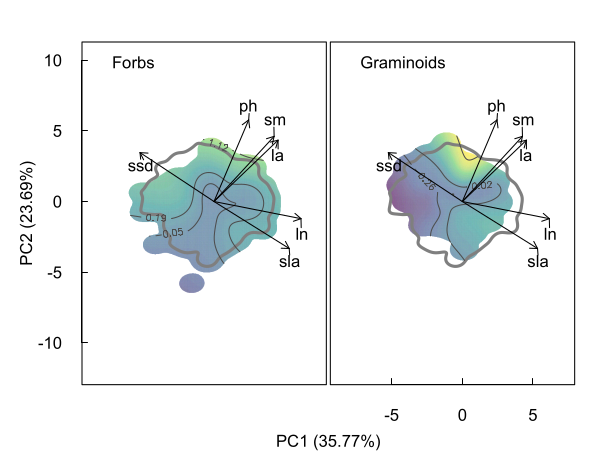

**Fig. S15. Mapping persistence in the aboveground trait space for forbs (left column) and graminoids (right column).** Details on the mapping can be found in **Appendix S2.** Trait abbreviations as in **Table S1**.

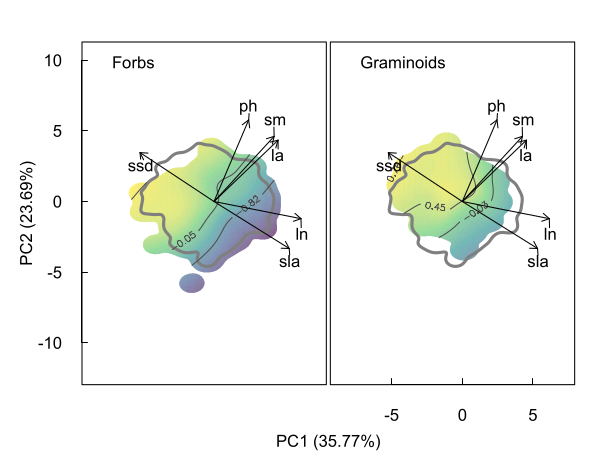

**Fig. S16. Mapping total bud-bank size in the fine-root trait space for forbs (left column) and graminoids (right column).** Details on the mapping can be found in **Appendix S2.** Trait abbreviations as in **Table S1**.

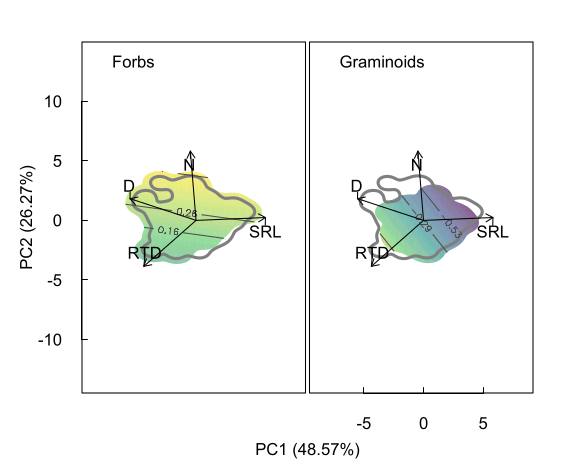

**Fig. S17. Mapping multiplication rate in the fine-root trait space for forbs (left column) and graminoids (right column).** Details on the mapping can be found in **Appendix S2.** Trait abbreviations as in **Table S1**.

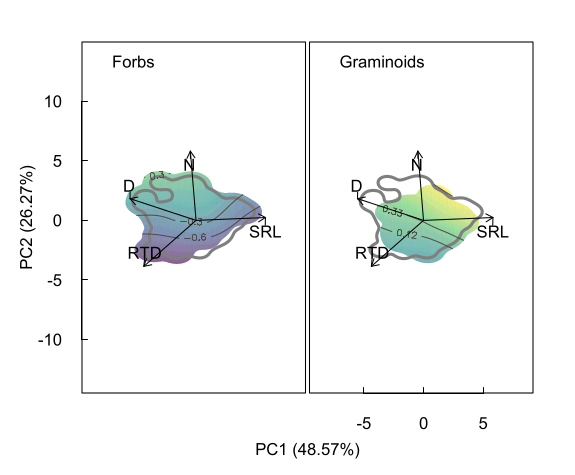

**Fig. S18. Mapping lateral spread in the fine-root trait space for forbs (left column) and graminoids (right column).** Details on the mapping can be found in **Appendix S2.** Trait abbreviations as in **Table S1**.

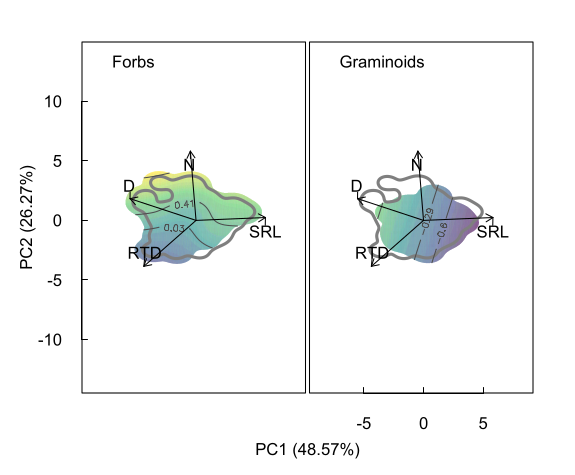

**Fig. S19. Mapping persistence in the fine-root trait space for forbs (left column) and graminoids (right column).** Details on the mapping can be found in **Appendix S2.** Trait abbreviations as in **Table S1**.

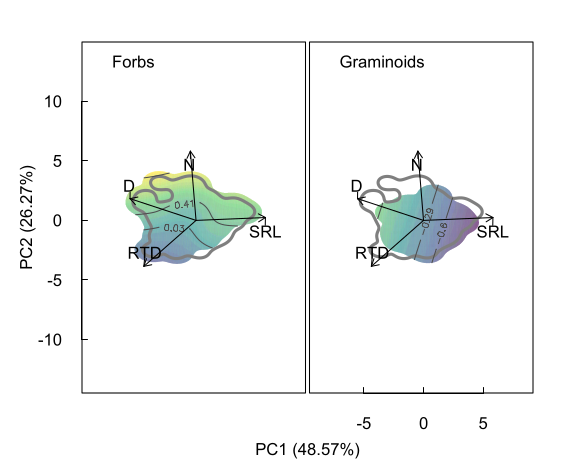

**Table S3. Full GAM statistics.** Statistics are reported for the models run on all data pooled and separately for forbs and graminoids (see main text for details) using either above or fine-root trait combinations as predictor, and each clonal-related trait as the response variable. Sample size (n), estimated degrees of freedom (Edf), the F-statistic and the p-value of the model are shown.

| Predictor | Response | Data | n | Edf | F-statistic | R^2^ | p-value |
| --- | --- | --- | --- | --- | --- | --- | --- |
| Aboveground | Bud-bank size | Pooled | 1257 | 14.96 | 5.71 | 0.08 | < 0.0001 |
| traits |  | Forbs | 919 | 14.86 | 7.5 | 0.11 | < 0.0001 |
|  |  | Graminoids | 338 | 12.33 | 3.01 | 0.13 | < 0.0001 |
|  | Multiplication rate | Pooled | 1215 | 13.86 | 3.31 | 0.05 | < 0.0001 |
|  |  | Forbs | 881 | 10.47 | 3.77 | 0.06 | < 0.0001 |
|  |  | Graminoids | 334 | 2.00 | 0.05 | 0 | 0.948 |
|  | Lateral spread | Pooled | 1216 | 11.7 | 2.79 | 0.04 | < 0.001 |
|  |  | Forbs | 880 | 11.53 | 2.07 | 0.04 | < 0.01 |
|  |  | Graminoids | 336 | 14.16 | 3.47 | 0.16 | < 0.0001 |
|  | Persistance | Pooled | 1224 | 11.85 | 21.48 | 0.22 | < 0.0001 |
|  |  | Forbs | 888 | 10.59 | 11.3 | 0.16 | < 0.0001 |
|  |  | Graminoids | 336 | 10.23 | 4.25 | 0.15 | < 0.0001 |
| Fine-root | Bud-bank size | Pooled | 195 | 2 | 2.04 | 0.01 | 0.13 |
| traits |  | Forbs | 110 | 2 | 0.15 | 0.01 | 0.86 |
|  |  | Graminoids | 85 | 2 | 3.08 | 0.05 | 0.05 |
|  | Multiplication rate | Pooled | 194 | 2 | 1.3 | 0.003 | 0.27 |
|  |  | Forbs | 110 | 3.38 | 1.17 | 0.03 | 0.32 |
|  |  | Graminoids | 84 | 2.58 | 0.9 | 0.01 | 0.45 |
|  | Lateral spread | Pooled | 195 | 2 | 6.87 | 0.06 | < 0.01 |
|  |  | Forbs | 110 | 4.25 | 1.34 | 0.06 | 0.25 |
|  |  | Graminoids | 85 | 2 | 3.32 | 0.05 | < 0.05 |
|  | Persistance | Pooled | 194 | 5.1 | 3.17 | 0.1 | < 0.01 |
|  |  | Forbs | 109 | 4.2 | 1.39 | 0.06 | 0.22 |
|  |  | Graminoids | 85 | 3.75 | 1.51 | 0.07 | 0.19 |
